# A Developmental Role for Microglial Presenilin 1 in Memory

**DOI:** 10.1101/2021.01.26.428181

**Authors:** Jose Henrique Ledo, Estefania P. Azevedo, Lucian Medrihan, Jia Cheng, Hernandez M. Silva, Kathryn McCabe, Michael Bamkole, Juan J. Lafaille, Jeffrey M. Friedman, Beth Stevens, Paul Greengard

**Affiliations:** Laboratory of Molecular and Cellular Neuroscience, The Rockefeller University, New York, New York, 10065, USA; Laboratory of Molecular Genetics. The Rockefeller University, New York, New York, 10065, USA; Skirball Institute of Biomolecular Medicine, New York University, New York, New York, 10016, USA; Department of Neurology, F.M. Kirby Neurobiology Center, Boston Children’s Hospital, Harvard Medical School, Boston, MA 02115, USA; Stanley Center for Psychiatric Research, Broad Institute of MIT and Harvard, Cambridge, MA, USA; Howard Hughes Medical Institute, Boston Children’s Hospital, Boston, MA., US

## Abstract

Microglia, the macrophages of the brain, are increasingly recognized to play a key role in synaptic plasticity and function; however, the underlying mechanisms remain elusive. Presenilin 1 (PS1) is an essential protein involved in learning and memory, through neuronal mechanisms. Loss of Presenilin function in neurons impairs synapse plasticity and causes cognitive deficits in mice. Surprisingly, here we show memory *enhancement* in mice by deleting PS1 selectively in microglia. We further demonstrate increased synapse transmission and *in vivo* neuronal activity in mice by depleting PS1 during microglial development, but not after microglial maturation. Remarkably, conditional deletion of PS1 in microglia during development increased memory retention in adulthood and was dependent on the NMDA receptor subunit GluN2B. In vivo calcium imaging of freely behaving mice revealed increased amplitude of neuronal Ca2+ transients in the CA1 hippocampus of *PS1 cKO* mice compared to control mice, suggesting a greater CA1 engagement during novel object exploration. Finally, loss of PS1 in microglia mitigated synaptic and cognitive deficits in a mouse model of Alzheimer’s disease. Together our results reveal a novel mechanism and function of PS1 in microglia in which modulation can enhance neuronal activity, learning and memory in mice.

## Introduction

Microglial cells are increasingly recognized to modulate neuronal function, including synaptic function and plasticity that could underlie learning and memory. [1–8]. Presenilin 1 (Psen1 gene – PS1 protein) is an essential protein for learning and memory [9–13] and it has been extensively studied in neurons in the context of Alzheimer’s disease (AD), as the most common genetic mutations known to cause familial Alzheimer’s disease (FAD) are found in PS1. Selective loss of Presenilin function in neurons impairs synapse plasticity and causes cognitive deficits in mice. These deficits were associated with reductions in synaptic levels of NMDA receptors and NMDA receptor-mediated responses [13]. Interestingly, PS1 is also expressed in microglia from development to adulthood [14, 15]; although the functional relevance is completely unknown.

In the adult mouse brain cortex, microglial cells show high expression of PS1, almost a 3-fold increase compared to neurons [15]. We previously reported that not only is PS1/γ-secretase active in microglia, but a gain of function mutant PS1 in microglia (PS1 ^KI/KI^) promotes synapse loss in the hippocampus of mice partly due to defective spine elimination by microglia [16]. Furthermore, we found that PS1 ^KI/KI^ alters microglial gene networks and number of progenitors during development [17], raising questions about the functional and behavioral consequences of PS1 loss of function in brain resident macrophages. Here, we hypothesized that loss of microglial PS1 function during development impacts neuronal plasticity and function in the adult brain.

To test this, we used different transgenic mouse models, including Cx3Cr1-Cre(ERT2) mice (for details see Supplemental Figure S1 A-J) and Tmem119-Cre(ERT2) mice to target brain microglia (for details see Supplemental Figure S7). We first assessed the effect of microglial *PS1 cKO* on hippocampal synaptic transmission using whole-cell patch clamp electrophysiology. Conditional knockout of PS1 in microglia led to increased frequency of miniature excitatory postsynaptic currents (mEPSC) compared to *control* mice (Figure 1 A-D). The mean decay tau but not the rise time of mEPSCs was also increased in CA1 pyramidal neurons from *PS1 cKO* mice (Figure 1 E-F). Our experiments also showed increased frequency in the spontaneous excitatory postsynaptic activity (sEPSC) of CA1 neurons of *PS1 cKO* mice compared to control littermates (Supplemental Figure S2 A-D).

**Figure 1:**
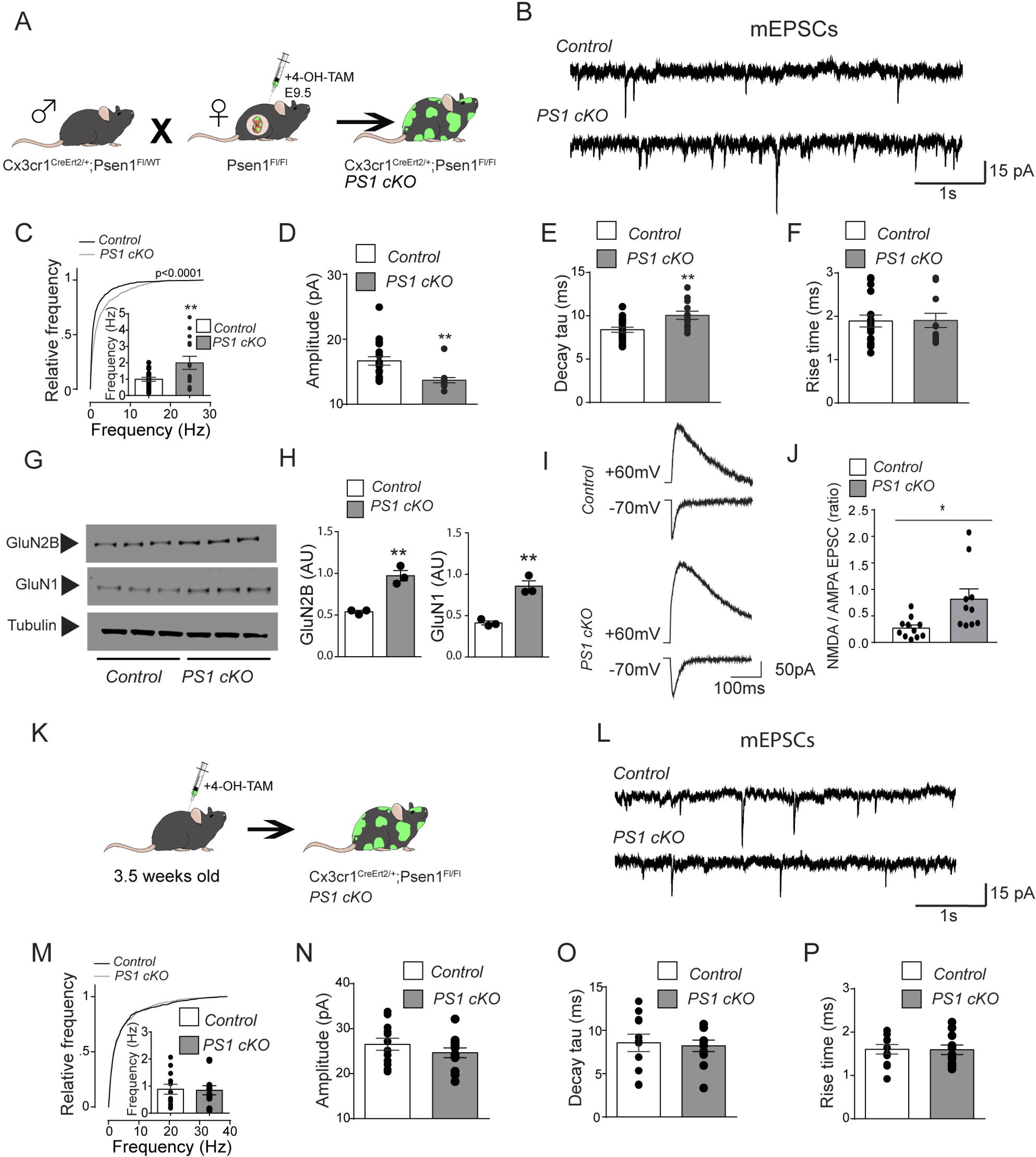
Increased NMDA receptor-mediated excitatory postsynaptic currents induced by absence of PS1 in developing microglia. (A) Schematic diagram showing the strategy to generate PS1 cKO mouse (Related to Figure 1 B–J). (B) Representative traces of mEPSCs in CA1 pyramidal neurons from acute hippocampal slices of *control* and *PS1 cKO* mice. (C) Mean cumulative probability of mEPSCs frequencies from *control* and *PS1 cKO.* Bin size for cumulative comparison was 2 Hz. Inlet histogram shows mean frequency is increased in *PS1 cKO* mouse compared to *control* mouse, each dot representing a recorded neuron; n= 19 neurons/5 mice for *control* and 14/4 for *PS1 cKO*. (D) Graph showing the mean amplitude of mEPSCs in CA1 pyramidal neurons from *PS1 cKO*. Each dot represents a recorded neuron; n= 19 neurons/5 mice for *control* and 14/4 for *PS1 cKO*. (E) Graphs showing that the mean decay tau but not the rise time (F) of mEPSCs is increased in CA1 pyramidal neurons from *PS1 cKO* mice. Each dot represents a recorded neuron; n= 19 neurons/5 mice for *control* and 14/4 for *PS1 cKO*. ** p < 0.01, two-tailed unpaired student *t* test (C and E) - Mann Whitney test (D and F). (G) Synaptosome-enriched fractions from *control* and *PS1 cKO* CA1 brains probed with indicated antibodies by Western blot. Lanes 1-2-3 represent *control*, lanes 4-5-6 represent *PS1 cKO.* (H) Densitometric quantification of the western blots **P < 0.01 using unpaired Student’s *t*-test. N=3 mice per group. Synaptosomes preparations from 2 months old mice. (I) Representative traces and (J) histogram showing mean of the NMDAR- to AMPAR-EPSC ratio of *control* versus *PS1 cKO* mouse. Scale bar, 50 pA, 100 ms. *P < 0.05 using unpaired Student’s *t*-test. N=10-11 neurons per group, 3 mice per group. Each graph bar represents ± SEM. (K) Schematic diagram showing the strategy to generate microglial PS1 knockout mouse after microglial maturation (Related to Figure 2 K – P). (L) Representative traces of miniature excitatory postsynaptic currents (mEPSCs) in CA1 pyramidal neurons from acute hippocampal slices of *control* and *PS1 cKO* mice (M) Mean cumulative probability of mEPSC frequencies from *control* and *PS1 cKO* mice. Bin size for cumulative comparison was 2 Hz. Inlet histogram shows mean frequency with each dot representing a recorded neuron; n= 11 neurons/4 mice for *control* and 12/4 for *PS1 cKO*. (N) Graph showing that mean amplitude of mEPSCs is unchanged in CA1 pyramidal neurons from *mmc-PS1 cKO* mice; n= 11 neurons/3 mice for *control* and 12/4 mice for *PS1 cKO*. (O) Graphs showing that the mean decay and (P) the rise time of mEPSCs is unchanged in CA1 pyramidal neurons from *control* and *PS1 cKO.* Each dot represents a recorded neuron; n= 11 neurons/4 mice for *control* and 12/4 for *PS1 cKO*. Each graph bar represents ± SEM, unpaired Student’s *t*-test, no statistically significant difference between groups (M-P).

To better understand the synaptic alterations after targeting PS1 in microglia, we examined the levels of various pre- and post-synaptic proteins in CA1 synaptosome-enriched brain extracts. We generated synaptosome-enriched fractions from *PS1 cKO* mouse brain and compared them to those from the *control* mice. First, we investigated the levels of the glutamate receptors subunits and found the NMDA receptors subunits GluN2B and GluN1 to be significantly increased in synaptosome-enriched preparations from *PS1 cKO* mouse brains (Figure 1 G-H). We found no differences in AMPA receptors subunits GluA1 and GluA2 levels (Supplemental Figure S3 A-B). In addition, no differences in the levels of vesicular glutamate transporter (VGlut), synaptophysin 1 and PSD95 levels of the *PS1 cKO* mice were found compared to *control* mice (Supplemental Figure S3 A-B).

We nexte examined the ratio of NMDA- to AMPA-EPSC, which provides a measure of NMDAR activity relative to the one of AMPAR regardless of the number of presynaptic afferents recruited and position of stimulating electrode. As shown in Figure 1I and J, the NMDAR- to AMPAR-EPSC ratio was significantly increased in CA1 pyramidal neurons from *PS1 cKO* mice compare to the age-matched controls (*control*: 0.27 ± 0.06, n = 11, cKO: 0.82 ± 0.19, n = 10; p < 0.05, *t*-test), suggesting that NMDAR activity is enhanced in the CA1 neurons of *PS1 cKO* mice. Interestingly, GluN2B overexpression has been shown to enhance memory in mice [18]. Importantly, when we depleted PS1 after 3.5 weeks postnatally (schematic diagram Figure 1 K) we did not observe alterations in synaptic transmission (sEPSC and mEPSC) in the PS1 cKO compared to *control* mice (Figure 1 K-P and Supplemental Figure S4 A-D), suggesting depleting PS1 during microglial development, but not after synaptic and circuit maturation has long term functional consequences on synaptic function and plasticity.

To test the functional consequences of microglial PS1 loss of function on neuronal function *in vivo*, we monitored neuronal activity *in vivo* in freely behaving mice using optic fiber photometry [19] (Figure 2A, left panel). Labeling of the CA1 neurons was achieved by injecting an adeno-associated virus (AAV) carrying the GCaMP6s construct into the CA1 hippocampus (Figures 2A, middle panel). We verified the implantation site of the optic fiber and expression of the GCaMP6s and above the CA1 area (Figure 2A, right panel). Peak analysis was performed to determine the number of events per minute (frequency of Ca2+ transients) [20] and area under the curve was plotted for each event (amplitude of Ca2+ transients). Consistent with the *in vitro* data, we observed increased neuronal activity analyzed as total frequency of events, assessed by measuring Ca2+ transients in the CA1 neurons of *PS1 cKO* mice, compared with their *control* littermates, in the basal state (Supplementary Figure S5 A-B) and during an open field task (Supplementary Figure S5 C-D).

**Figure 2:**
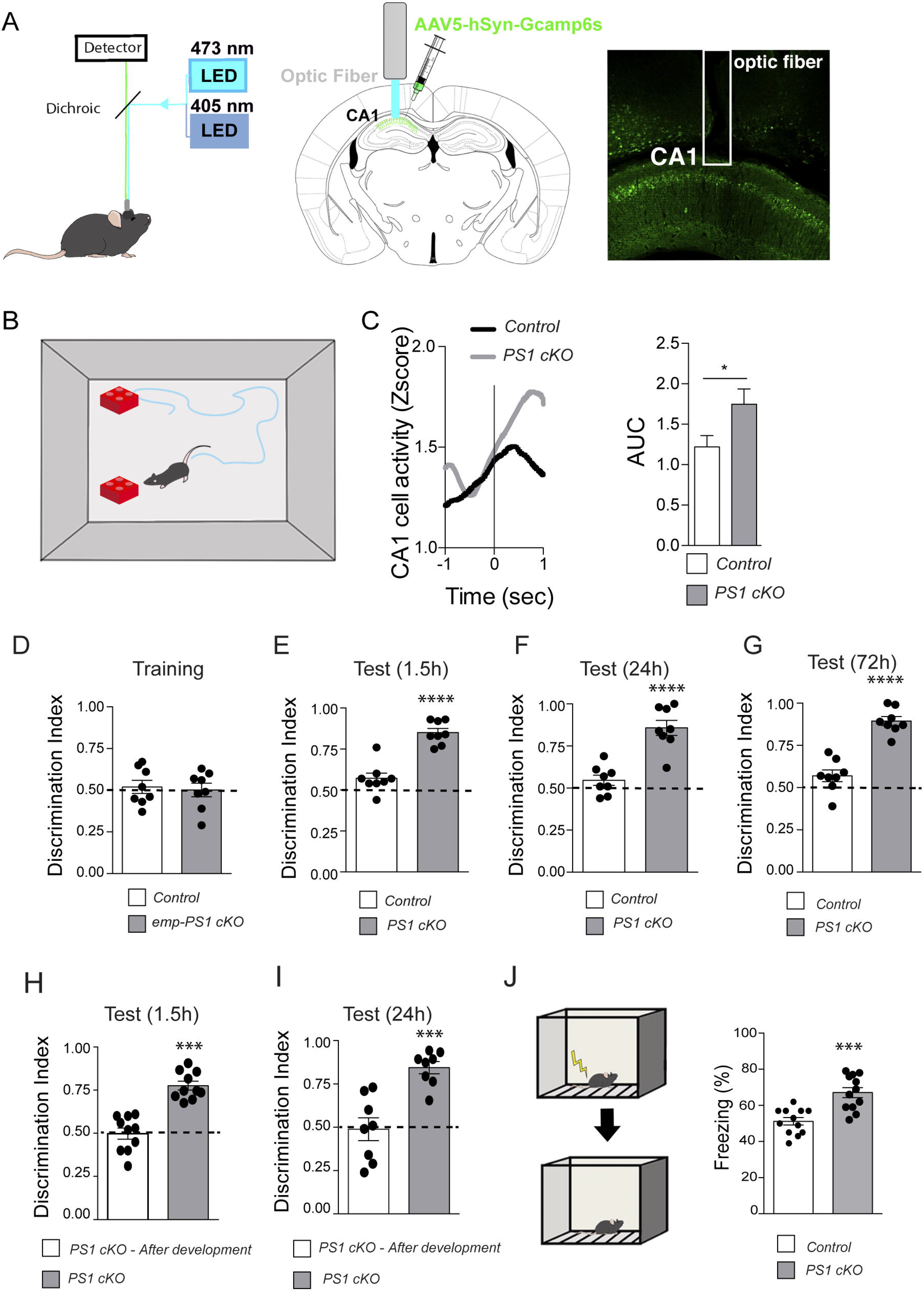
Enhanced neuronal activity and memory in mice lacking microglial PS1 during development. (A) Left panel, experimental scheme representing fiber placement in the mouse brain. Mice were attached to a fiber photometry setup (Doric) through patch cords and GCaMP6s was excited with a 473 nm LED (green) and a 405 nm LED (blue). Middle and Right panels, experimental scheme and representative images showing AAV5-hSyn-GCaMP6s expression in mouse CA1 hippocampus. (B) Neuronal activity was recorded during novel object exploration. Middle, average GCaMP6s z-score (black line, *control* and red line, *PS1 cKO*) across all recording sites time-locked to noseto-object touch. (C) Graph showing area under the curve (AUC) measurement for each nose-to-object time-locked Ca2+ transient for each animal trial; n=6 mice per group. (D) Exploratory preference in the training session. Discrimination index 0.5 (dashed lines) represents performance at chance (50%). The amount of time spent exploring both objects was the same for all mice. (E) Exploratory preference in *control* and *PS1 cKO* mouse at 1.5 h hour retention test, (F) 24h retention test and (G) 72h retention test. (H) Exploratory preference in *PS1 cKO* - *after development* (*PS1 cKO* in 3.5 weeks old mouse, white bar) or *PS1 cKO* (PS1 cKO in embryonic development, gray bar) at 1.5h retention test and (I) 24h retention test. Mice were 2-3 months old. We used a 3-min training protocol (see Methods) as this does not typically lead to changes in short-term memory or long-term memory in wild-type mice. Data are represented as mean ± SEM, n ≥ 6-10 mice per group. *P < 0.05, ***P < 0.001, ****P < 0.0001 using unpaired Student’s *t*-test. (J) Mice were trained in contextual fear conditioning and tested 24h after training. Compared with *control* mice, *PS1 cKO* resulted in increased freezing. Data are represented as mean ± SEM, n= 12 mice per group. **P < 0.01, unpaired Student’s *t*-test.

Next, we asked whether the PS1 cKO during development altered neuronal activity when mice were engaged in an object recognition task that requires CA1 neurons [21]. To analyze the dynamics of CA1 neurons during memory encoding, we recorded the neuronal activity from CA1 neurons in freely moving mice during the training phase of novel object recognition task (Figure 2B). When neuronal activity recordings were time-locked with behavior, in this case nose-to-object exploration, CA1 neuronal activity was increased (Figure 2C). Interestingly, the amplitude of Ca2+ transient was increased in *PS1 cKO* mice compared to mice, suggesting a greater CA1 engagement during novel object exploration (Figure 2C).

To determine whether increased neuronal activity during novel object exploration affects the cognitive state of the mice, we monitored the performance of *control* and *PS1 cKO* mice in a memory task. To evaluate memory performance, we measured memory retention in the novel object recognition task. In this test, memory retention was evaluated after 1.5 h, 24h, 72h (for details see Methods) after a 3 min training session as this does not typically lead to changes in short-term memory or long-term memory in wild-type mice [22–24]. We reasoned that by observing improved performance of the tasks after a brief period of training it would indicate that there was memory enhancement in the *PS1 cKO* mice. During the training both groups did not show preference for a specific object (Figure 2D). To test memory retention, one of the familiar objects used in the training session was replaced by a novel object. The animals were allowed to explore both objects for 5 min and the time spent exploring the new and familiar object was expressed as discrimination index (0.5 represents performance at chance 50%, for details see methods). At 1.5-h, 24h or 72h retention test, the *PS1 cKO* mice but not the wild-type mice showed an increased discrimination index for the novel object, indicating that those mice retained the memory of the old object longer (Figure 2 E-G). According to our data, when we depleted PS1 after 3.5 weeks postnatally, the performance of the *PS1 cKO* mice at 1.5h or 24h was similar to wild type (Figure 2 H-I, white bars).

We next assessed associative emotional memory in these mice through contextual fear conditioning (Figure 2J). Results revealed increased freezing in *PS1 cKO* mice compared to *control* mice at the 1 d retention delay (Figure 2J). Importantly, *PS1 cKO* mice did not show motor alterations or anxiety-like behavior compared to *control* mice as revealed by open field test (Supplementary Figure S6 A-E). Together, our data showed that CA1 neurons activity increase during object exploration, especially in the *PS1 cKO* mice, suggesting the engagement of these neurons during the memorization of object-place location. The greater activity of these neurons in the *PS1 cKO* mice also correlate with the increased memory retention but only when PS1 is depleted during microglial development, not after microglial maturation.

In order to further validate our findings and test that *PS1 cKO* increased GluN2B levels and increase memory retention through microglia but not non-microglial brain macrophages (NMBM), we used the novel *Tmem119 ^CreErt2/+^* mice line, which provides the currently most specific targeting of microglia. Importantly, *Tmem119 ^CreErt2/+^; Psen1 ^Fl/Fl^* (*PS1 cKO*) mice showed increased GluN2B levels and improved memory performance in the novel object recognition test compared to *Tmem119 ^CreErt2/+^* (*control*) mice (Supplemental Figure S7 A-D), consistent with our finding in *Cx3cr1 ^CreErt2/+^* mice (Figure 1G and 2D-G), further supporting the conclusion that the cKO of PS1 in microglia increases memory retention in mice.

We next asked whether conditional deletion of PS1 in microglia is beneficial in the context of a disease model that exhibits memory and plasticity defects. Given the established genetic risk associated with PS1 mutation and Alzheimer’s disease, and our findings that PS1 deletion enhances learning and memory in healthy mice, we next tested whether deletion of PS1 in microglia could mitigate synaptic and memory deficits in an established AD mouse model. Recognizing that mouse models of AD do not recapitulate the full spectrum of Alzheimer’s, these mice nonetheless provide a valuable tool to study particular memory impairment. For instance, memory impairment is well-established in the Alzheimer’s mouse model J20-PDGF-APPSw,Ind (hAPP) [25].

To investigate whether a cKO of PS1 in microglia could also enhance neuronal activity and thus, memory in a mouse model of memory losswe crossed the *hAPP* mice with the *PS1 cKO* mice (*hAPP *PS1 cKO)* and evaluated CA1 neuronal activity in hAPP and hAPP * *PS1 cKO* mice using in vivo fiber photometry. Consistent with the results from *PS1 cKO* mice without the hAPP transgene, we found an increased number of events per minute in CA1 neurons in *hAPP *PS1 cKO* mice compared with their *control* littermates at baseline (Supplementary Figure S8 A-B) or during open field task (Supplementary Figure S8 C-D). The difference in amplitude between hAPP and hAPP * *PS1 cKO* mice were also greater when mice were engaged in spatial exploration, both during open field (Supplementary Figure S8 C-D) or novel object exploration during a memory task (Figure 3 A and B). These data also suggest that CA1-specific circuits are malfunctioning in hAPP mice during tasks that require spatial navigation.

**Figure 3:**
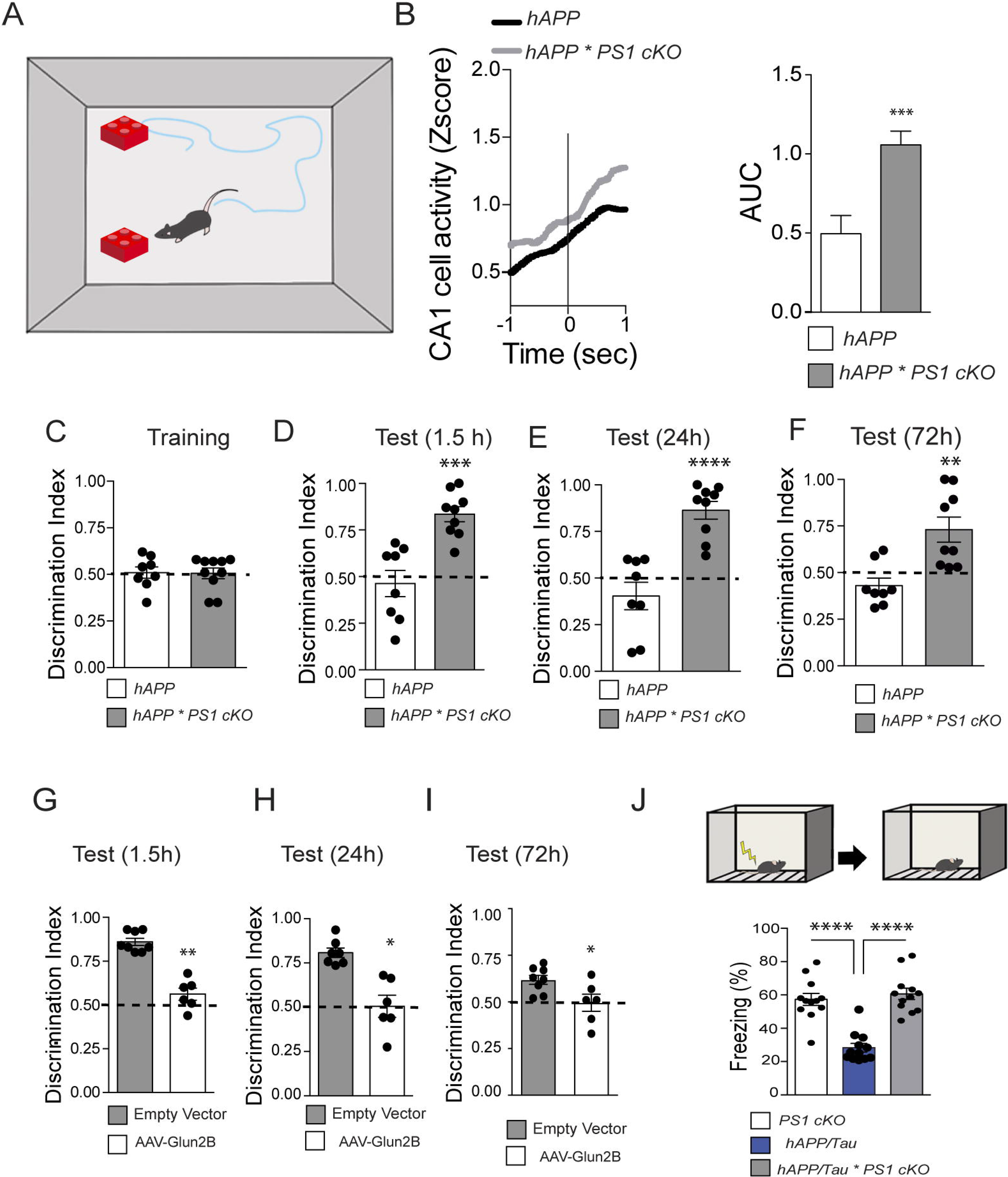
Enhanced neuronal activity and memory in Alzheimer’s disease mouse models induced by PS1 cKO. (A) Neuronal activity was recorded during object exploration. (B) Average GCaMP6s z-score (black line, hAPP and red line, *hAPP* * *PS1 cKO*) across all recording sites time-locked to nose-to-object touch. (B) Graphs showing area under the curve (AUC) measurement for each nose-to-object time-locked Ca2+ transient for each animal trial; n=6. Data are represented as mean ± SEM. ***P < 0.001 using unpaired Student’s t-test. (C) Exploratory preference in the training session. Discrimination index 0.5 represents performance at chance (50%). The amount of time spent exploring both objects was the same for all mice. (D) Enhanced exploratory preference in *hAPP* * *PS1 cKO* mouse at 1.5-hour retention test, (E) 24h retention test and (F) 72h retention test. (G) Exploratory preference in *PS1 cKO* mouse, injected with an AAV-CRISPR-Cas9 system (all-in-one SaCas9 and sgRNA, white bar) targeting Glun2B or *control* (AAV vector empty, gray bar), at 1.5h retention test, (H) 24h retention test and (I) 72h retention test. Mice were 5-6 months old. We used a 3-min training protocol (see Methods) as this does not typically lead to changes in short-term memory or long-term memory in wild-type mice. Data are represented as mean ± SEM, n ≥ 6-10 mice per group. *P < 0.05, **P < 0.01, ***P < 0.001, ****P < 0.0001 using unpaired Student’s *t*-test. (J) Mice were trained in contextual fear conditioning and tested 24h after training. Compared to AD mouse, *PS1 cKO* resulted in increased mouse freezing. Data are represented as mean ± SEM, n =12 mice per group. Data were analyzed using ANOVA, post hoc (Bonferroni). ****P < 0.0001.

To test whether *PS1 cKO* could rescue memory deficits in AD mouse, we monitored the performance of the *hAPP* and *hAPP *PS1 cKO* mice in the novel object recognition task. No differences were observed between mouse groups during the training session (Figure 3 C). However, at 1.5-h, the *hAPP *PS1 cKO* mice exhibited a preference towards the novel object, indicating that they were able to retain the memory of the old object, and recognize the new object as being novel, after 1.5 hour (Figure 3 D, gray bar). On the other hand, the *hAPP* mice failed to show preference towards the novel object, indicating that they were not able to retain the memory of the old object after 1.5 hour (Figure 3 D, white bar). Similarly, at 24h or 72h (Figure 3 E-F), only the *hAPP *PS1 cKO* mice exhibited preference for the novel object, confirming that the *PS1 cKO* improved memory retention in *hAPP* AD mice.

To test whether the memory enhancement in *PS1 cKO* mice was mediated by GluN2B. We knocked down GluN2B using an AAV vector to deliver a CRISPR-Cas9 system (SaCas9 and sgRNA, AAV-CRISPR) in the CA1 hippocampus of the *PS1 cKO* mice (Supplemental Figure S9). We performed the novel object recognition task in *hAPP * PS1 cKO* mouse injected with AAV-CRISPR *control* (empty vector, gray bars) or AAV-CRISPR targeting GluN2B (white bars) (Figure 3 G-I). Our results reveal that at 1.5h, 24h and 72h retention test *hAPP * PS1 cKO* mice were not able to retain the memory of the old object when injected with AAV-CRISPR targeting Glun2B (graph white bars), but they were able to retain the memory of the old object when injected with AAV-CRISPR *control* (empty vector, graph gray bars) (Figure 3 G-I). These results indicate that microglial *PS1 cKO* increased memory retention through GluN2B.

We next assessed associative emotional memory in *PS1 cKO* and AD mice through the contextual fear conditioning. Due to the fact that contextual fear conditioning impairment in the J20-PDGF-APPSw,Ind mice is controversial [27, 28], we next performed fear conditioning test in the APPSwe,tauP301L mouse (48 weeks old) to assess associative emotional memory in *control,* APPSwe,tauP301L and *PS1 cKO* crossed to APPSwe,tauP301L (APPSwe,tauP301L* *PS1 cKO*) mice. Results revealed that APPSwe,tauP301L mice showed a significant decreased freezing at the 1 d retention delay compared to the *PS1 cKO* mice (Figure 4J). On the other hand, the APPSwe,tauP301L* *PS1 cKO* showed significant increased freezing compared to APPSwe,tauP301 mouse (Figure 3J). Our study presents evidence for a novel role of microglial PS1 during development that ultimately affects cognition.

The precise mechanism by which loss of PS1 in microglia increases GluN2B in neurons remains to be elucidated. NMDA receptors play a central role in the development of neural circuits. Interestingly, NMDA receptor subunits composition changes during forebrain development [29]. This shift in NMDA receptors composition is highly evident in the first 3 weeks in the developing brain [29, 30]. Thus, it will be important to investigate the mechanisms by which the cKO of PS1 in microglia increased GluN2B in that developmental time window. Data showed that developmental changes in GluN2B levels occur independent of transcriptional or translational changes [30]. Furthermore, Zhang, et al., (2013) found that NMDA receptors are required for pruning and strengthening of immature synapses [31].

Selective preservation and strengthening of GluN2B-dominated synapses might be part of the mechanism involved in the GluN2B activity induced by PS1 cKO. Intriguingly, Hayashi et al., (2006) found that microglia-conditioned medium (MCM) potentiated NMDA receptor-mediated synaptic currents. This effect was suggested to be mediated by the NMDA receptor co-agonist glycine secreted by microglia [32]. Whether glycine secreted by microglia is involved in the NMDA-dependent effects of microglial PS1 cKO needs further investigation. Finally, it will be important to investigate in future studies whether increased memory retention induced by PS1 cKO is due to the lack of γ-secretase activity and whether it involves γ-secretase substrates TREM2 and SIRPa. Overall, our study opens a new avenue for the investigation of microglia-induced memory retention in health and disease.

## Author Contribution

J.H.L., conceived; J.H.L., B.S. and P.G., supervised this study; J.H.L. designed experiments; J.H.L., E.P.A., L.M., K.M. H.MS. performed and analyzed experiments; J.H.L., prepared figures. J.H.L. and B.S. wrote the paper.

## Acknowledgments

This work was supported by funds received from Fisher Center for Alzheimer’s Research Foundation, JPB Foundation, and Cure Alzheimer’s Fund. J.H.L. is a Pew Latin American Fellow in the Biomedical Sciences, supported by The Pew Charitable Trusts.

