## Supplementary figures and images for "A Developmental Role for Microglial Presenilin 1 in Memory"

### Supplemental Figure S1

#### Supplemental Figure 1

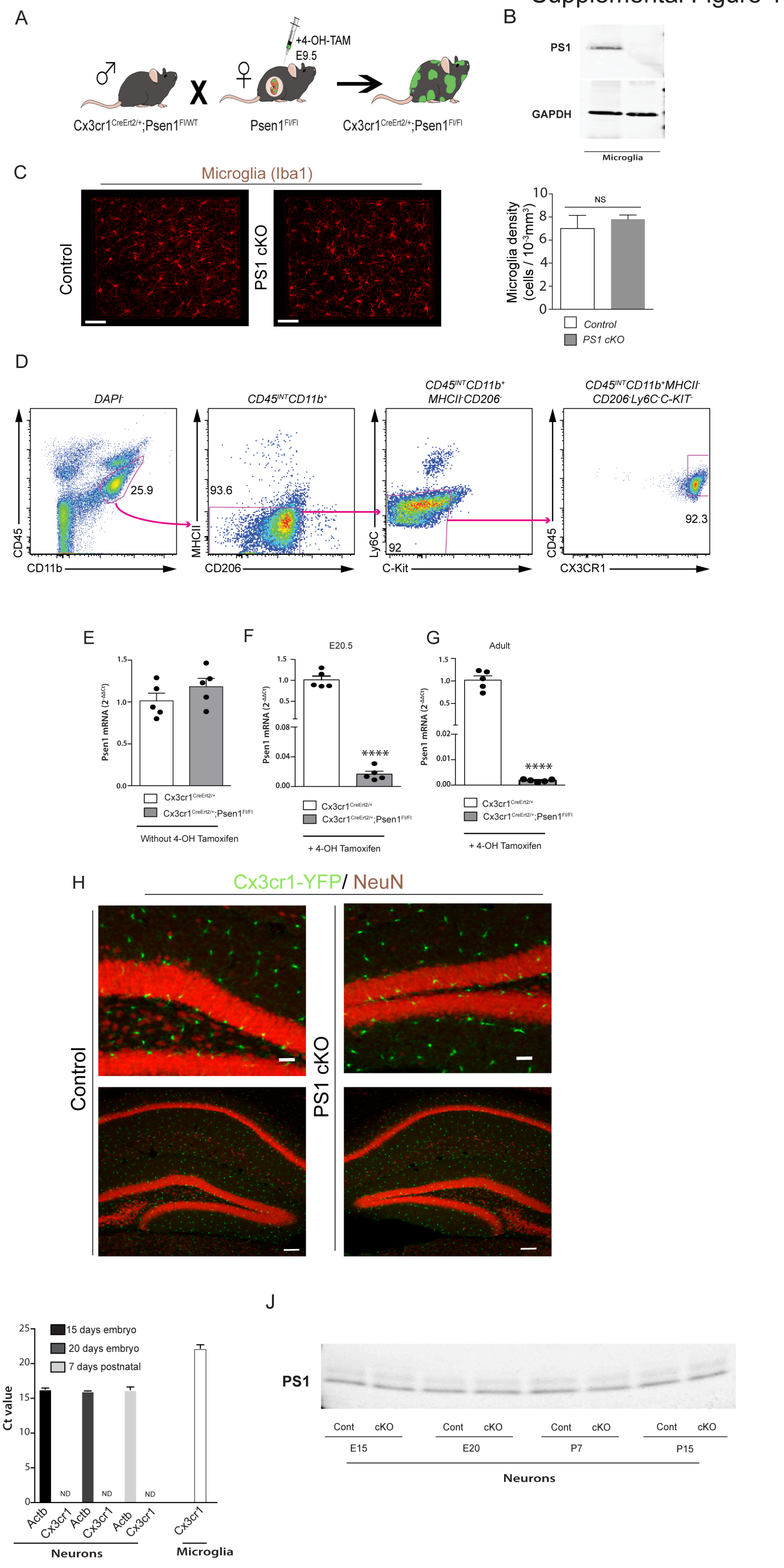

### Supplemental Figure S2

#### Supplemental Figure 2

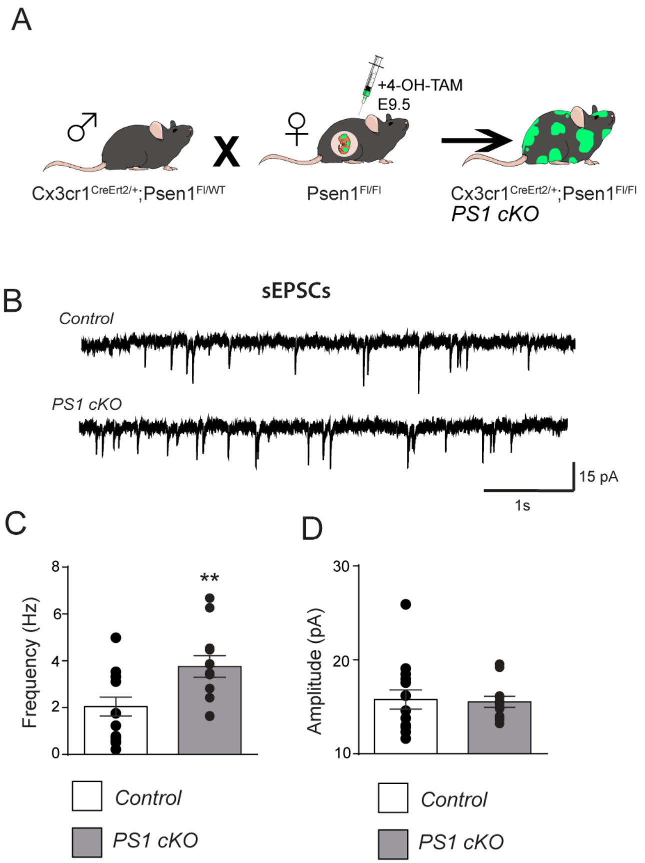

### Supplemental Figure S3

#### Supplemental Figure 3

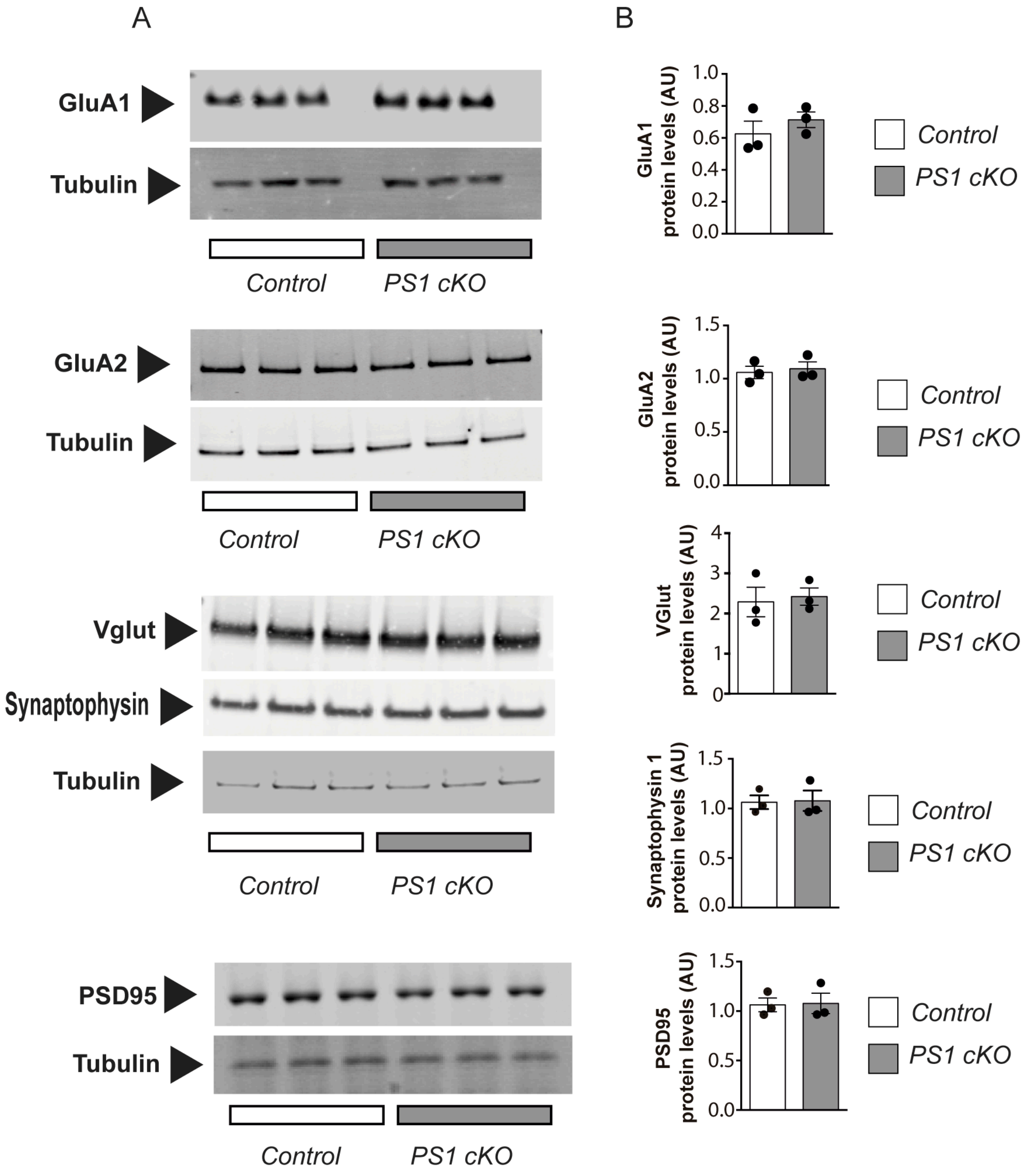

### Supplemental Figure S4

#### Supplemental Figure 4

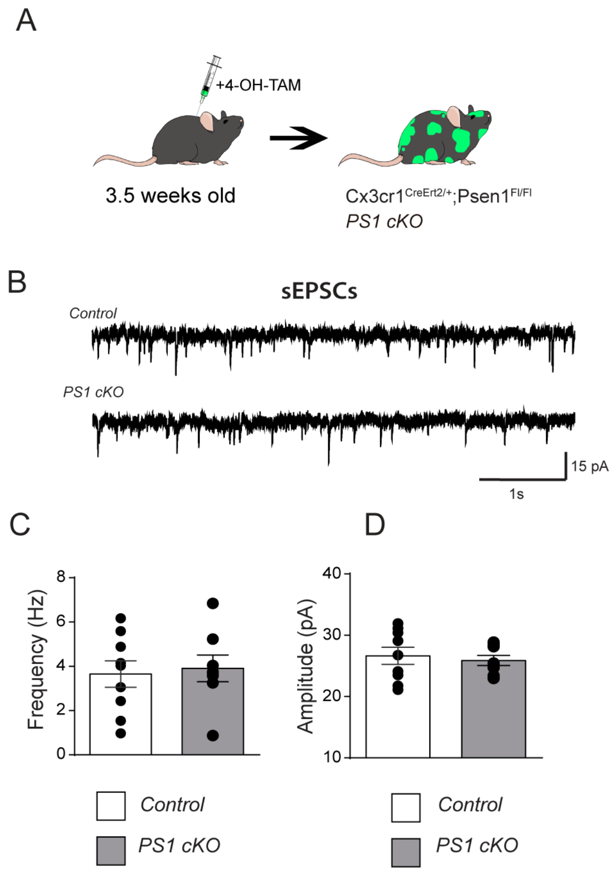

### Supplemental Figure S5

#### Supplemental Figure 5

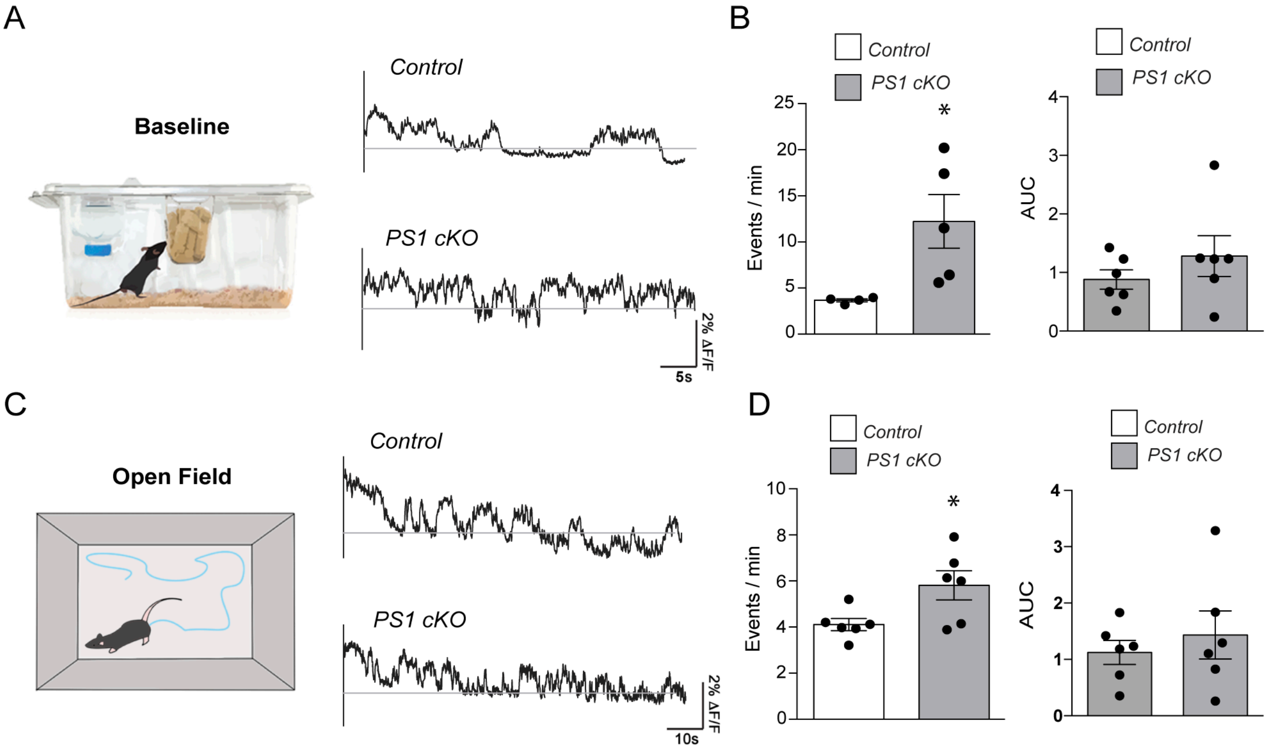

### Supplemental Figure S6

#### Supplemental Figure 6

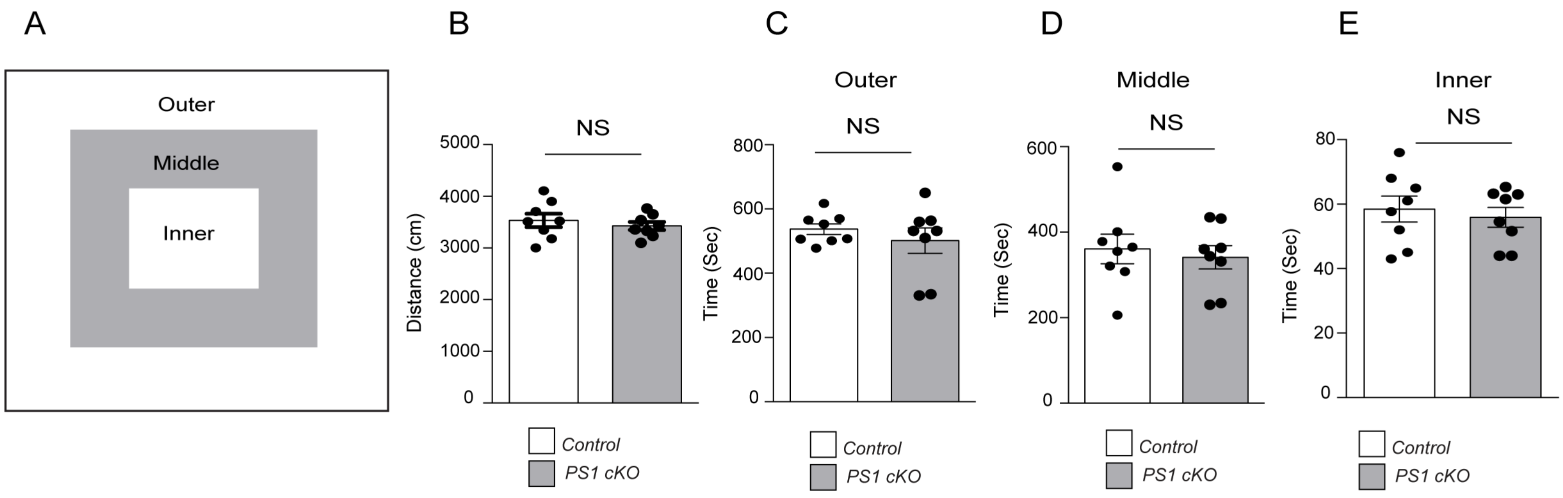

### Supplemental Figure S7

#### Supplemental Figure 7

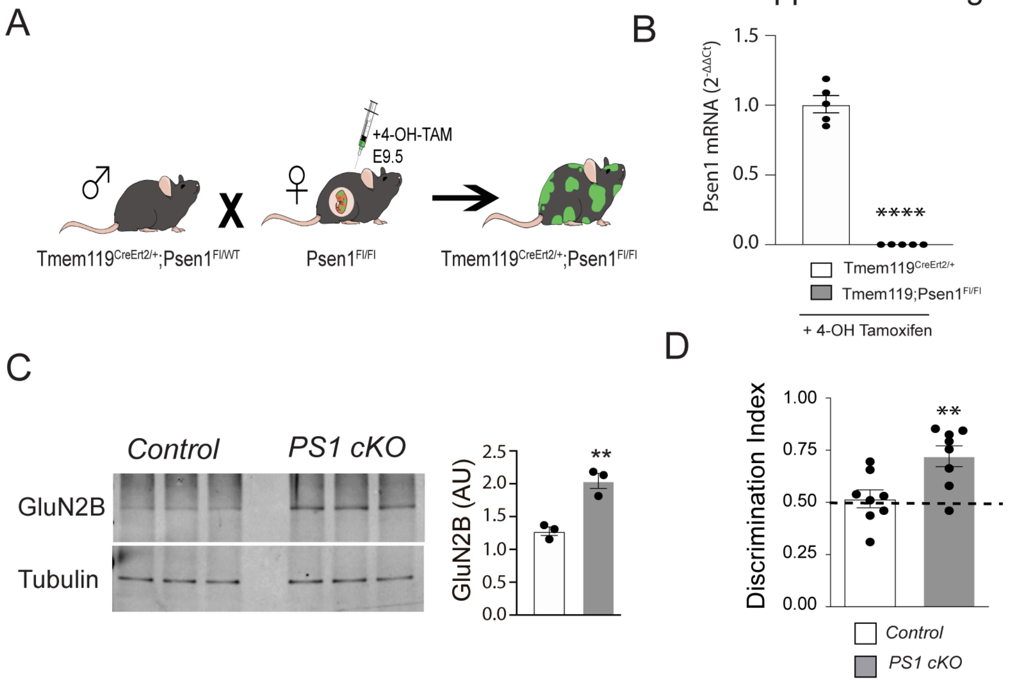

### Supplemental Figure S8

#### Supplemental Figure 8

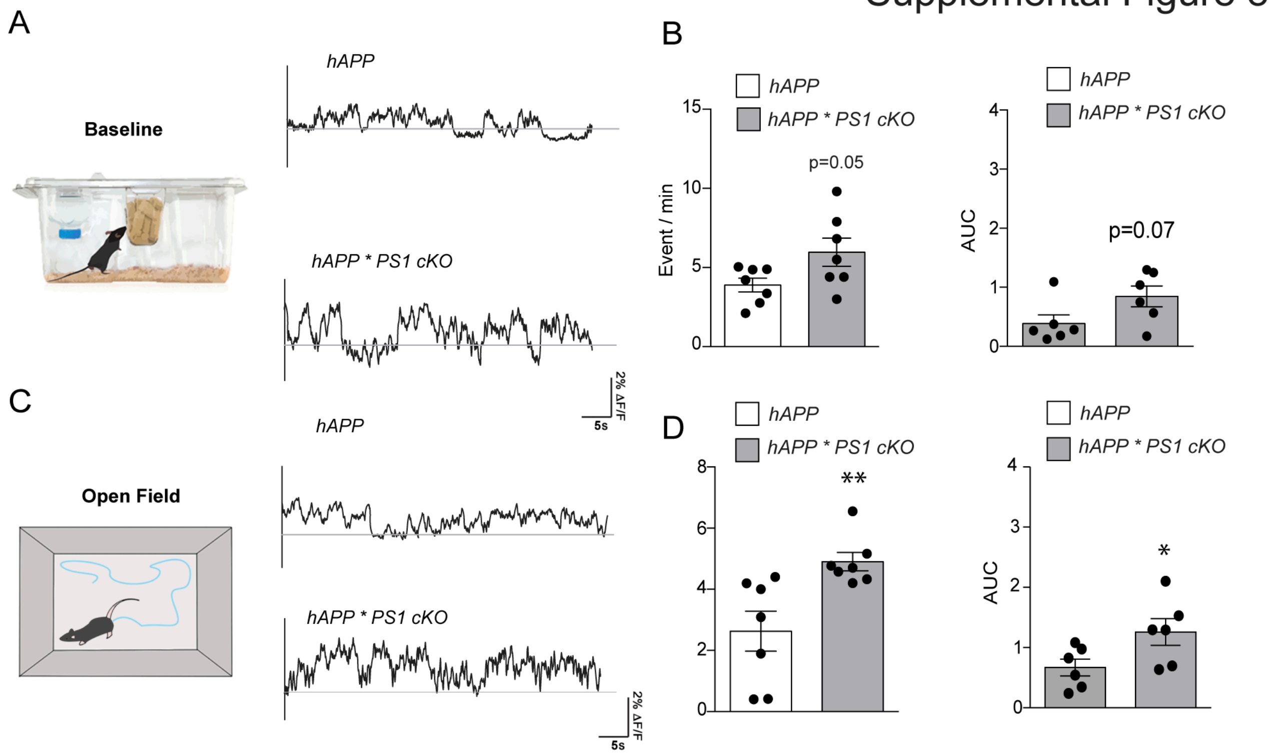

### Supplemental Figure S9

Supplemental Figure 9

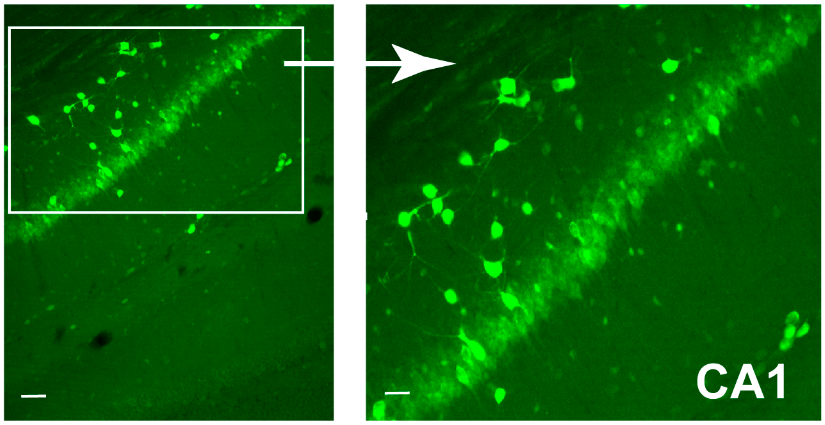
