## Supplemental Text for "A Developmental Role for Microglial Presenilin 1 in Memory"

### Supplemental Figures Legends

#### Supplemental Figure S1: PS1 deletion in microglia.

(A) Schematic diagram showing the strategy to generate *PS1 cKO* mouse. (B) PS1 depletion in microglia sorted from *control* (*Cx3Cr1*$^{CreErt2/+}$) and *PS1 cKO* (*Cx3Cr1*$^{CreErt2/+}$; *Psen1* $^{Fl/Fl}$) mice and probed with PS1 antibody by Western blot. (C) Confocal stack of microglia in the 2-month-old *control* (*Cx3Cr1*$^{CreErt2/+}$) and *PS1 cKO* (*Cx3Cr1*$^{CreErt2/+}$; *Psen1* $^{Fl/Fl}$) mouse brain. Scale bar represents 50 µm. Quantification of microglial density in *control* and *PS1 cKO* mice (cells / $10^{-3}$ $mm^3$). Data represent means ± SEM (*n* = 5 mice per group). (D) Gating strategy to sort microglia. Brains were harvested and gated for $CD45^{INT}$, $CD11B^+$, $CX3CR1^+$, $CSF1R^+$, C-$KIT^-$, LY-$6C^-$, $CD206^-$, $MHCII^-$. (E) Quantitative RT-PCR of *Psen1* on microglia from *Cx3Cr1*$^{CreErt2/+}$ and *Cx3Cr1*$^{CreErt2/+}$; *Psen1* $^{Fl/Fl}$ adult mouse prior to 4-OH tamoxifen treatment. (F) Quantitative RT-PCR of *Psen1* on microglia from *Cx3Cr1*$^{CreErt2/+}$ and *Cx3Cr1*$^{CreErt2/+}$; *Psen1* $^{Fl/Fl}$ embryos after to 4-OH tamoxifen treatment. (G) Quantitative RT-PCR of *Psen1* on microglia from *Cx3Cr1*$^{CreErt2/+}$ and *Cx3Cr1*$^{CreErt2/+}$; *Psen1* $^{Fl/Fl}$ adult mouse after to 4-OH tamoxifen treatment. (H) Coronal sections from *control* (*Cx3Cr1*$^{CreErt2/+}$) and *PS1 cKO* (*Cx3Cr1*$^{CreErt2/+}$; *Psen1* $^{Fl/Fl}$) mouse stained with anti-Cx3cr1 (green) for microglia and NeuN (red) for neurons. Representative z-stack images with maximum projection are shown. Data are represented as mean ± SEM. Two-tailed t-test, ****P < 0.0001, N = 5 mice or embryos per group. Scale bar, 50 µm (right panels). (I) Quantitative RT-PCR of *Actb* (Actin, beta) and *Cx3cr1* on neuronal cells isolated from *Cx3Cr1*$^{CreErt2/+}$ mice at E15.5, E20.5 and P7 and microglial culture (white bar). ND = non detectable, Cx3cr1 expression was not detected in neuronal cells. (J) Neuronal cells isolated from *control* and *PS1 cKO* CA1 brains probed with PS1 antibody by Western blot at day E15.5, E20.5, P7 and P15.

**Supplemental Figure S2: Absence of PS1 in microglia during development increased spontaneous excitatory postsynaptic currents.**

(A) Schematic diagram showing the strategy to PS1 knockout in microglial development. (B) Representative traces of spontaneous excitatory postsynaptic currents (sEPSCs) in CA1 pyramidal neurons from acute hippocampal slices of *control* and *emp-PS1 cKO* mice. (C) Histograms showing that mean frequency but not amplitude (D) of sEPSCs is increased in CA1 neurons from *PS1 cKO* mice (frequency: 2.04 ± 0.4 Hz in *control* vs 3.75 ± 0.4 Hz in *PS1 cKO* mice; amplitude: 15.77 ± 0.9 pA in *control* vs 15.52 ± 0.5 pA in *PS1 cKO* mice; n= 14 neurons/5 mice for *control* and 12/4 mice for *PS1 cKO*. Each dot represents a recorded neuron. ** $p < 0.01$, two-tailed unpaired student *t* test.

**Supplemental Figure S3: Western blot analysis of synaptic proteins from CA1 hippocampus of PS1 cKO and Control mice.**

(A) Synaptosome-enriched fractions from *Control* and *PS cKO* CA1 brains probed with indicated antibodies by Western blot. Lanes 1-2-3 represent *Control*, lanes 4-5-6 represent *PS1 cKO*. (B) Densitometric quantification of Western blots in (A) **$P < 0.01$ using unpaired Student's t-test. N=3 mice per group, bar graphs represent mean ± SEM.

**Supplemental Figure S4: Absence of PS1 in microglia after microglial maturation does not increased spontaneous excitatory postsynaptic currents.**

(A) Schematic diagram showing the strategy to PS1 knockout mouse after microglial maturation. (B) Representative traces of spontaneous excitatory postsynaptic currents (sEPSCs) in CA1 pyramidal neurons from acute hippocampal slices of *control* and *PS1 cKO* mice. (C) Histograms showing that both mean frequency and (D) amplitude of sEPSCs is unchanged in CA1 pyramidal neurons from *PS1 cKO* mice compared to littermate controls (frequency: 3.64 ± 0.6 Hz in *Control* vs 3.9 ± 0.6 Hz in *PS1 cKO*; amplitude: 26.65 ± 1.4 pA in *control* vs 25.89 ± 0.8 pA in *PS1 cKO*; n= 9 neurons/3 mice for *control* and 8/3 mice for *mmc-PS1 cKO*). Each dot represents a recorded neuron. ** p < 0.01, two-tailed unpaired student *t* test.

**Supplemental Figure S5: PS1 cKO enhanced neuronal activity during baseline and open field exploration.**

(A) Neuronal activity recording at baseline or (C) during open field test were performed for 5 min in Control and PS1 cKO mouse. (A and C, right panels) Representative Ca2+ traces (represented as the percent change in fluorescence over the mean fluorescence [ΔF/F]) for each group is shown in the middle panels. Peak analysis of Ca2+ traces. PS1 cKO in increased events / min (B and D, left panel). Histogram showing area under the curve (AUC) measurement for each animal trial (B and D, right panel). Data are represented as mean ± SEM. N=4-6 mice per group *P < 0.05 using unpaired student's t-test.

**Supplemental Figure S6: PS1 cKO does not cause anxiety-like behavior in mice.**

Increase excitability of CA1 neurons in *PS1 cK*O mouse does not impact basal anxiety-like behavior assessed in the open field test. Statistically significant differences were not detected between the *control* and *PS1 cKO* mice in total distance traveled (B) time spent in outer (C), middle (D) and inner zone (E) of the arena. Data are represented as mean ± SEM, n= 8 mice per group.

**Supplemental Figure S7: Increased GluN2B levels and enhanced novel-object recognition memory in *Tmem119* $^{CreErt2/+}$; *Psen1* $^{Fl/Fl}$ (*PS1 cKO*) mice.**

(A) Schematic diagram showing the strategy to generate *PS1 cKO* mice using *Tmem119* $^{CreErt2/+}$ mice. (B) Quantitative RT-PCR of *Psen1* on microglia from *Tmem119* $^{CreErt2/+}$ and *Tmem119* $^{CreErt2/+}$; *Psen1* $^{Fl/Fl}$ adult mouse after 4-OH tamoxifen treatment. Data are represented as mean ±

SEM, n =5 mice per group. ****P < 0.0001, using unpaired Student's *t*-test. (C) PS1 depletion in microglia. Microglia from *Tmem119* $^{CreErt2/+}$ (*control*) and *Tmem119* $^{CreErt2/+}$; *Psen1* $^{Fl/Fl}$ (*PS1 cKO*) probed with indicated antibodies by Western blot. Lanes 1-2-3 represent *control*, lanes 4-5-6 represent *PS1 cKO*. Data are represented as mean ± SEM, n =3 mice per group. **P < 0.01, using unpaired Student's *t*-test. (D) Novel Object Recognition test. Discrimination index 0.5 (dashed lines) represents performance at chance (50%). Exploratory preference in *control* and *PS1 cKO* mouse at 24 hour retention test. In this test, memory retention was evaluated 24h after a 3 min training session as this does not typically lead to changes in short-term memory or long-term memory in wild-type mice (for details see Methods). Data are represented as mean ± SEM, n =8 mice per group. **P < 0.01, using unpaired Student's *t*-test.

**Supplemental Figure S8: PS1 cKO enhanced neuronal activity in an Alzheimer's disease mouse model during open field exploration.**

(A) Neuronal activity recording at baseline or (C) during open field test were performed for 5 min in hAPP and hAPP **PS1 cKO* mice. (A and C, right panels) Representative Ca2+ traces (represented as the percent change in fluorescence over the mean fluorescence [ΔF/F]) for each group is shown in the middle panels. Peak analysis of Ca2+ traces, PS1 cKO increased events / min (B and D left panels). Histogram showing area under the curve (AUC) measurement for each animal trial (B and D, right panel). Data are represented as mean ± SEM. N=6-7 mice per group. *P < 0.05, **P < 0.01 using unpaired student's t-test.

**Supplemental Figure S9: AAV-mediated expression of SaCas9/CRISPR systems to target GluN2B in mouse brain.**

Representative images showing AADJ-MCP292597-AD01-3-A00 expression (green) in mouse CA1 hippocampus. Scale bars, 100 ⌠m (left panel) and 55 ⌠m (right panel).

### METHODS

#### Mice

C57BL/6 (000664), Cx3Cr1$^{CreErt2/+}$ (021160), Psen1$^{loxP}$ (004825), hAPP - PDGF-APPSw/Ind (34836-JAX) mice were purchased from the Jackson Laboratories and maintained in our facilities. APPSwe,tauP301L -1Lfa Psen1$^{tm1Mpm}$ (34830-JAX) mouse was crossed to WT mouse. Mice APPSwe,tauP301L (hemizygous) were used in the experiments. Presenilin 1 deletion: Psen1$^{loxP/loxP}$ mice have been described previously (*1*). Psen1$^{loxP/loxP}$ (homozygous) and CX3CR1$^{CreER/+}$ (heterozygous) mice were purchased from the Jackson Laboratory (stock no. 004825) and (stock no. 021160) respectively. Psen1 $^{loxP/loxP}$ (homozygous) mice were first crossed to CX3CR1$^{CreER/+}$ (heterozygous). Psen1 knockout mice were generated by further crossing CX3CR1$^{CreER/+}$ : Psen1 $^{loxP/+}$ to Psen1 $^{loxP/loxP}$ (homozygous). CX3CR1$^{CreER/+}$ (heterozygous) mice littermates were used as controls. Mice were weaned at the third postnatal week, genotyped by Transetyx using real-time PCR and kept on a 12 h/12 h light/dark cycle (lights on at 7:00) with access to food and water ad libitum. Mice were maintained at The Rockefeller University Animal facilities and used at 10-12 weeks of age for all experiments except when otherwise indicated. Littermates of the same sex were randomly assigned to experimental groups. Both female and male mice were used for experiments. Animal care and experimentation were according to NIH guidelines and were approved by the Institutional Animal Care and Use Committee at The Rockefeller University (protocol #18035-H).

#### 4-Hydroxytamoxifen treatment

For Cre-dependent recombination during microglial development - 8 weeks old pregnant female mouse received two doses of 0.4 g/g of 4-hydorxytamoxifen at embryonic day E9.5 and E11.5.

For Cre-dependent recombination after microglial maturation - 3.5 weeks old mouse received two doses of 0.4 g/g of 4-hydorxytamoxifen, injections were 2 days separately. Specific target of PS1 after tamoxifen induced Cre recombination was accessed (Supplemental Figure S1).

**Brain microglia isolation from adult mice**

Mice were perfused with DPBS (Ca2+/Mg2+-free) and brains were placed in FACS buffer (PBS, 5 % FBS and 10 mM HEPES). Brain were minced and incubated with 4000 U/mL of collagenase D (Roche, 11088858001) at 37 °C for 30 min. Collagenase was inactivated by adding 10 mM EDTA for 5 min at 37 °C. Digested brains were passed through a 70-µm cell strainer, washed in FACS buffer and submitted to centrifugation at 2,000 r.p.m. in 38% Percoll for 30 min. Cell pellets were resuspended in FACS buffer and incubated with a CD16- and CD32-specific antibody (BD-Pharmingen 553141) for 15 min to ensure that nonspecific binding to FC receptors was blocked. Cells were then washed and stained with the markers described below to certify cell population specificity

**Flow cytometry analysis**

Fluorescent-dye-conjugated antibodies were purchased from Biolegend (anti-CD115 (Csfr1), 135510; anti-F4/80, 123131; anti-Cx3Cr1, 149016; anti-CD117 (c-kit), 105824; anti-Ly6c, 128033; anti-P2RY12, 848004), Invitrogen (Anti-CD11b, 47-0112-82; anti-CD45, 56-0451-82) or BD-Pharmingen (anti-CD16/CD32 (FC blocking); 553141). Live cells were verified using DAPI (4',6-Diamidino-2-Phenylindole, Dilactate), (D3571, Thermofisher). Flow cytometry data were acquired on an LSR-II flow cytometer (Becton Dickinson) and analyzed using FlowJo software (Tree Star).

#### Real-time relative quantification PCR

Microglial cells isolated from mice brain (see above) were sorted directly into RNA lysis buffer (Qiagen, 79216) supplemented with 2 M dithiothreitol (DTT). RNA was isolated as described above. qPCR was performed using Taqman reagents. ActB were used to normalize samples. Predesigned probes used were purchased from IDTDNA. mRNA levels are expressed using the $2^{-\Delta\Delta Ct}$ method (*2*).

#### Purification of Neurons

Neurons were purified with MACS technology. Neuronal Tissue Dissociation Kit (P) (Miltenyi Biotec 130-092-628) was used for dissociating mouse adult brain. For neuron-specific separation, Neuron Isolation Kit was used according to the manufacturer's protocol (Miltenyi Biotec 130-115-390).

#### Immunoblotting and antibodies

Mouse CA1 hippocampus were lysed with a Syn-PER Synaptic Protein Extraction Reagent (Syn-PER™, 87793, Thermo Fisher Scientific) supplemented with a protease and phosphatase inhibitor cocktail (78442, Thermo Fisher Scientific). Synaptosomes-enriched extracts were prepared according to the instructions provided by the manufacturer (see above). Protein levels were measured by the BCA method. The samples were mixed with the standard protein sample buffer and subjected to SDS-PAGE with 4–20% Novex Tris-Glycine gels (Thermo Fisher Scientific), followed by protein transfer onto a PVDF membrane. Immunoblotting was performed with a standard protocol using the following antibodies: anti-GluN2B (mouse monoclonal,75-097, NeuroMab, 1:1,000), anti-GluN1 (mouse monoclonal,75-272, NeuroMab, 1:1,000), anti-GluA1 (mouse monoclonal, 75-327, NeuroMab, 1:1000), anti-GluA2 (rabbit monoclonal, ab133477,

Abcam, 1:1000), anti-Synaptophysin 1 (Guinea pig polyclonal, 101004, Synaptic Systems, 1:1,000), anti-Vglut (mouse monoclonal, 135011, Synaptic Systems, 1:1000), anti-PSD95 (rabbit polyclonal, ab18258, Abcam, 1:1,000), anti- beta-Tubulin (rabbit polyclonal, 2146, Cell Signaling, 1:1,000), anti-GAPDH (goat polyclonal, GTX89740, GeneTex, 1:1000), Presenilin 1 (mouse monoclonal, MAB5232, Millipore Sigma, 1:500). Secondary antibodies were from Li-COR: IRDye® 800CW Donkey anti-Mouse (926-32212, 1:1000) IRDye® 800CW Donkey anti-Rabbit (926-32213, 1:1000), IRDye® 680RD Donkey anti-Rabbit (926-68073, 1:1000).

**Electrophysiology**

Mice were euthanized with CO2. After removal of the brains, transversal slices (400 μm thickness) were cut using a Vibratome 1000 Plus (Leica Microsystems, USA) at 2 °C in a cutting solution containing (in mM): 87 NaCl, 25 $NaHCO_3$, 2.5 KCl, 0.5 $CaCl_2$, 7 $MgCl_2$, 25 glucose, and 75 sucrose saturated with 95% $O_2$ and 5% $CO_2$. Slices were left to recover for 45 minutes at 35 °C and next, for 1 h at RT in the recording solution (see below). The extracellular solution used for recordings contained (in mM): 125 NaCl, 25 $NaHCO_3$, 2.5 KCl, 1.25 $NaH_2PO_4$, 2 $CaCl_2$, 1 $MgCl_2$ and 25 glucose (bubbled with 95% $O_2$ and 5% $CO_2$). The slice was placed in a recording chamber (RC-27L, Warner Instruments, USA) and constantly perfused with oxygenated aCSF at 24 °C (TC-324B, Warner Instruments, USA) at a rate of 1.5–2.0 ml/min. CA1 pyramidal neurons were selected for recording based on their shape and position in the CA1 layer using an upright Olympus BX51WI microscope (Olympus, Japan). Whole-cell patch-clamp recordings were performed with a Multiclamp 700B/Digidata1550A system (Molecular Devices, Sunnyvale CA, USA) and glass pipettes (King Precision Glass, Inc, Glass type 8250) pulled in a horizontal pipette puller (Narishige) to a resistance of 3–4 MΩ. The intracellular solution contained (in mM): 126 Kgluconate, 4 NaCl, 1 $MgSO_4$, 0.02 $CaCl_2$, 0.1 BAPTA, 15 glucose, 5 HEPES, 3 ATP, 0.1 GTP (pH 7.3). Recordings of spontaneous excitatory postsynaptic currents (sEPSCs) were performed in the presence of 30μM bicuculline to block GABA activity. For recordings of miniature excitatory postsynaptic currents (mEPSCs), tetradotoxin (0.5 μM) was added in the extracellular solution to block $Na^+$ currents. Data were acquired at a sampling frequency of 50 kHz and filtered at 1 kHz and analyzed offline using pClamp10 software (Molecular Devices, Sunnyvale, CA, USA). All electrophysiological data are expressed as means ± SEM. Statistical analysis was performed using the two-tailed unpaired Student's *t*-test or Mann-Whitney test (GraphPad Prism 5) as designated in figure legends.

Whole-cell voltage-clamp recording technique was performed to record evoked EPSCs as described previously (*5*). Dorsal CA1 hippocampal slices (400 μm) were positioned in a perfusion chamber attached to the fixed stage of an upright microscope and submerged in continuously flowing oxygenated ACSF. Bicuculline (20 μM) were added in the ACSF to block GABA receptors. Patch electrodes contained internal solution (in mM): 130 Cs-methanesulfonate, 10 CsCl, 4 NaCl, 10 HEPES, 1 MgCl2, 5 EGTA, 2 QX -314, 12 phosphocreatine, 5 MgATP, 0.2 Na3GTP, pH 7.2-7.3, 265-270 mOsm. CA1 pyramidal neurons were visualized with a 40X water-immersion lens and recorded with the Multiclamp 700A amplifier (Molecular Devices). EPSC was evoked by stimulation of the Schaffer-collateral fibers by a bipolar stimulating electrode (FHC). Stimulation pulse was generated from a stimulation isolation unit controlled by a S48 pulse generator (Grass Technologies). Evoked AMPAR-EPSC (eAMPA-EPSC) was elicited by a stimulation pulse (0.5 ms, 70 μA) when cells were clamped at -70 mV. For evoked NMDAR-EPSC, cells were first clamped at -70 mV and then depolarized to +60 mV for 3 s to fully relieve the voltage-dependent Mg2+ block. Same stimulation pulse was then applied to elicit EPSC. eAMPA-EPSC was calculated as the peak amplitude. eNMDA-EPSC was calculated at 100ms after stimulation. Data analyses were performed with Clampfit, and GraphPad Prism 6.

**Stereotaxic injections and Optic Fiber Implantation**

Mice were anesthetized with 2% isoflurane, placed in a stereotaxic frame (Kopft Instruments). Eye ointment was applied to the eyes and a subcutaneous injection of meloxicam (2 mg/kg) was given to each mouse after surgery for up to 2 days. Hair was shaved, the scalp was disinfected with iodine solution and an incision was made. The craniotomy was performed using a dental drill (Dremel). AAV was infused using a hamilton syringe at a rate of 0.1 ⌠L/min at a total volume of 0.3 ⌠L. After viral infusion, the needle was kept at the injection site for 10 min and then slowly withdrawn. High titer ($10^{11}$) viral preparations of AAV5-hSyn-GCaMP6s were purchased from Addgene and injected unilaterally at the following coordinates: from bregma, AP: -1.82 mm, ML:0.5 mm, DV:-1.7 mm. For all surgeries, mice were monitored for 72h to ensure full recovery and 2-4 weeks later mice were used in experiments. Optic fiber implants (400 ⌠m, Doric) for photometry experiments were implanted to the same coordinates as the viral injections with a -0.2 mm difference. Implants were inserted slowly and secured to the mouse skull using two layers of Metabond (Parkell Inc). The Mice were single housed in flat top cages and monitored in the first weeks following optic implantation. AAV-Crispr to target GluN2B was purchased from GeneCopoeia (AADJ-MCP292597-AD01-3-A00: AAVPrime™ Purified AAV Particles titer >/=$5x10^{12}$ GC/ml CRISPR all-in-one (SaCas9 and sgRNA), 3 x target sites Grin2b; AADJ-CCPCTR01-AD01-100: sgRNA Scrambled Control AAV Particles).

### Optic Fiber photometry

Mice were acclimated to patch cables for 5 min for 3 days before experiments. Analysis of the signal was done using fiber photometry system and processor (RZ5P) from TDT (Tucker-Davis Technologies), which includes the Synapse software (Tucker-Davis Technologies). Post-recording analysis was performed using custom-written MATLAB codes. The bulk fluorescent signals from each channel were normalized to compare across animals and experimental sessions. The 405 nm LED emission was used as the isosbestic control. GCaMP6s signals that are recorded at this wavelength are not calcium-dependent; thus, changes in signal can be attributed to autofluorescence, bleaching, and fiber bending. Accordingly, any fluctuations that occurred in the 405 control channels were removed from the 473 channel before analysis. Change in fluorescence (ΔF) was calculated as (473 nm signal − fitted 405 nm signal), adjusted so that a ΔF/F was calculated by dividing each point in ΔF by the 405 nm curve at that time. Z-score and plot traces were calculated for all experiments using MATLAB. Peak analysis to determine frequency and amplitude of the signals was done by determining the median average deviation (MAD) of the corrected/normalized data set and peaks were identified as events that exceeded the MAD by 2.91 deviations (*6*). Behavioral variables (nose-to-object touch during object exploration) were timestamped in the signaling traces via the real-time processor as TTL signals from Noldus (Ethovision) software. This allowed for precise determinations of the temporal profile of signals in relation to specific behaviors. All behavior was manually scored by a blind, second investigator who did not conduct the photometry experiment.

**Novel-object recognition (NOR) memory**

The NOR task was conducted as described in (*7*, *8*). For the NOR task, animals were acclimated to the behavior room where the task was held and handled for 3 min each day for at least 4 days before the task. Mice were habituated for 5 min in a 28 x 28 cm open field arena previously cleaned with 70% ethanol. During the training phase, a 3 min training session was used as this does not typically lead to changes in short-term memory or long-term memory in wildtype mice (*8–10*). Briefly, two identical objects are place in opposite sides of the arena and mice are free to explore the arena and objects for 3 min. No preference for a specific object was observed during the training session. After 1.5 h, 24h and 72h after the training session, mice performed the testing session of the task; in which one of the objects (familiar object) is replaced by a novel object and the mice were free to explore the arena and objects for 5 min. For training and test condition a discrimination index was calculated as the ratio between the time spent exploring the novel object and the sum of time spent exploring the familiar object and novel object. Arena and object were cleaned with 70% alcohol in between subjects. A distinct group of mice (*control vs PS1 cKO*) was used for each time point.

**Contextual Fear Conditioning**

We used a fear conditioning shock chamber (MedAssociates) coupled with shock-grid floors (Coulbourn Instruments). During training (5 min), mice were placed in the chamber and a foot shock was delivered (0.7 mA for 2s). After, mice were allowed to stay in the chamber for 1 min and then removed to their home cage. During the retention test, each mouse was placed back into the shock chamber and the freezing response was recorded for 5 min (contextual conditioning). Subsequently, the mice were put into a novel chamber and monitored for 5 min and freezing responses were also recorded. Freezing (i.e., absence of movement except for breathing) was measured manually by a blind second experimenter.

**Quantification and statistical analysis**

Statistical analysis was performed in GraphPad Prism or MATLAB software, in particular cases pClamp10 software (electrophysiology) was used. Optic fiber photometry raw data was recorded using TDT Synapse software (Tucker-Davis Technologies) and analyzed using custom written codes in MATLAB software. Animal behavior was time-locked with Ca2+ recording using digital timestamps Noldus Ethovision 9.0 software. Normality tests were performed before choosing the statistical test.
